# Perceived timing of active head movement at different speeds

**DOI:** 10.1101/258590

**Authors:** Carolin Sachgau, William Chung, Michael Barnett-Cowan

## Abstract

The central nervous system must determine which sensory events occur at the same time. Actively moving the head corresponds with large changes in the relationship between the observer and the environment, sensorimotor processing, and spatiotemporal perception. Numerous studies have shown that head movement onset must precede the onset of other sensory events in order to be perceived as simultaneous, indicating that head movement perception is slow. Active head movement perception has been shown to be slower than passive head movement perception and dependent on head movement velocity, where participants who move their head faster than other participants require the head to move even earlier than comparison stimuli to be perceived as simultaneous. These results suggest that head movement perception is slower (i.e., suppressed) when the head moves faster. The present study used a within-subjects design to measure the point of subjective simultaneity (PSS) between active head movement speeds and a comparison sound stimulus. Our results clearly show that i) head movement perception is *faster* when the head moves faster within-subjects, ii) active head movement onset must still precede the onset of other sensory events (Average PSS: -123 to -52 ms) in order to be perceived as occurring simultaneously even at the fastest speeds (Average peak velocity: 76°/s to 257°/s). We conclude that head movement perception is slow, but that this delay is minimized with increased speed. While we do not provide evidence against sensory suppression, which requires active versus passive head movement comparison, our results do rule out velocity-based suppression.

## 1. Introduction

To create an accurate representation of the world, the central nervous system (CNS) processes incoming signals from different sensory modalities and determines how the information from these senses relate to one another. The ability to bind sensory information accurately in time is crucial for the CNS to make correct decisions about our environment and our movements in it. Since the same event can stimulate multiple sensory modalities at different relative times, the CNS must distinguish whether these stimuli originated from the same or separate events. Actively moving the head corresponds with large changes in the relationship between the observer and the environment, sensorimotor processing, and spatiotemporal perception. While quickly detecting the onset of head movement is crucial for reflexive behaviour and rapidly updating the representation of the world around us, past research suggests that perceptual awareness of active head movement onset is slower than passive movement of the head, as well as slower than comparison stimuli such as light, touch or sound [1].

The vestibular system is essential for functions ranging from the perception of self-motion and spatial orientation, to the motor coordination for maintaining balance and posture [2]. The physiological response to vestibular stimulation is extremely fast. For example, the vestibulo-ocular reflex (VOR), which is the compensatory movement of the eyes in response to head movement, responds to vestibular stimulation in 5-6ms [3]. One may reasonably assume that the perceived timing of vestibular stimulation is equally fast. Surprisingly, research has shown that the perception of vestibular stimuli is slow compared to other senses [4; See 5 for a review]. One explanation for such a delay is that it may take time for the *perceptual* mechanisms of the CNS to accumulate enough evidence to determine that head movement onset has been initiated. For example, the types of computations needed to generate reflexes such as the VOR are relatively simplistic compared to the additional time required to determine position and velocity from acceleration through integration, the latter of which is the signal encoded by neurons at the vestibular periphery [6].

Barnett-Cowan and Harris [1] found that increased active head movement velocities resulted in greater perceptual delays, suggesting that the velocity of active head movements affect the ability to detect the onset of the movement. One theory attributed this to the suppression of the vestibular afferent signals found during higher movement velocities as recorded in monkeys [7], the rate of which was similar to the modulation of the point of subjective simultaneity (PSS) between the onset of active head movement and a comparison light, touch or sound stimulus for human temporal perception [1]. It was suggested that this suppression could stem from the reafference of the efference copy. Here, when a motor command signal is generated, a copy of the signal is created to attenuate the afferent information resulting from the motion. However, one issue with the findings of Barnett-Cowan and Harris [1] was that the results were from between-subjects data. Thus, their result that greater head movement velocities result in greater perceptual delays should really be interpreted as participants who move their head faster than other participants require the head to move even earlier than comparison stimuli to be perceived as simultaneous.

Indeed, there is good reason to suspect that within-subjects the perceived delay for head movement onset should be reduced as the head moves faster. Here, previous literature has shown that increased stimulus intensity reduces the delay in the perceived timing of that sense. Most studies that have investigated the effect of stimulus intensity on the perceived timing of sensory stimuli involve audiovisual tasks or comparing two visual events. As early as 1933, Smith [9] reported that stimuli of higher intensity were perceived earlier than lower intensity, in an audiovisual temporal order judgment (TOJ) task where the intensity of stimuli was varied. Roufs [10] showed that bright flashes of light are perceived earlier than synchronous dim flashes. When two flashes were shown simultaneously with different intensities between 10-1000 trolands, observers reported an apparent movement of the flash in the direction of the dimmer flash, due to the longer perceptual delay of the weaker flash. Efron [11] paired a light stimulus with a shock stimulus under four sets of conditions, where either stimuli could be weak or strong. If both stimuli were strong, there was less of a deviation from true simultaneity than if both stimuli were weak. Additionally, if either stimulus was weak, the weaker stimulus had to be presented before the stronger stimulus in order for the observer to subjectively rate them as occurring simultaneously. Neumann and colleagues [12] varied stimulus intensity in an audiovisual task, where for most trials the auditory stimulus had to be presented first in order to be perceived as simultaneous. This effect could be reversed, however, when the intensity of light was decreased, and the intensity of sound was increased. These results suggest that intensity can influence the order in which stimuli from different modalities are perceived. More recent studies confirm that higher intensity stimuli in audiovisual tasks are perceived earlier in time [13], and that higher intensity stimuli are less likely to be reported as synchronous than lower intensity stimuli in simultaneity judgement tasks [14]. With respect to the vestibular system, the only study we are aware of that has used a vestibular task, found that the PSS between the onset of passive self-motion and sound is significantly shorter during passive whole-body rotations when the rotation frequency increases from 0.5 Hz to 1 Hz (˜170 ms difference) and as the angular velocity increases from 5 to 60 °/s (˜133 ms difference)[15]. Taken together, these findings suggest that a greater velocity (stimulus intensity) should result in less time required for the head to move prior to other stimuli to be perceived as simultaneous.

In this study, we vary the velocity of an active head movement and analyze the data both between- as well as within-subjects. Participants performed TOJ tasks for slow, medium, and fast active head movements paired with an auditory stimulus, using a within-subjects design. After this, velocities for each individual were stratified into four conditions, to increase the amount of data points for a within-subjects analysis. We have three main hypotheses. First, to replicate the finding from previous studies [1, 4, 16,17; See 5 for a review] that an active head movement must precede a paired auditory stimulus for them to be perceived as simultaneous. Second, to replicate the findings from Barnett-Cowan and Harris [1] that an increase in active head movement velocity paired with an auditory stimulus will lead to a larger negative PSS between-subjects. Third, to explicitly test the relationship between active head movement velocity and PSS using a within-subjects design and predict, based on [1], that an increase in active head movement velocity paired with an auditory stimulus will lead to a larger negative PSS between-subjects.

## 2. Methods

### 2.1 Participants

20 participants (19-25y) who reported having no auditory, visual or vestibular disorders were remunerated $10 for one hour of testing. This study was carried out in accordance with the recommendations of Canada’s Tri-Council Policy Statement: Ethical Conduct for Research Involving Humans (TCPS2) by the University of Waterloo’s Human Research Ethics Committee with written informed consent from all subjects. All participants gave written informed consent in accordance with the Declaration of Helsinki.

### 2.2 Apparatus

Head movement was measured using the YEI 3-Space Sensor: Data-logging inertial measurement unit by Yost labs, which was mounted onto the back of the head using an elastic strap. The YEI 3-Space Sensor consists of a triaxial gyroscope, accelerometer, and compass. The measurements were recorded at 1000Hz using the Python API available directly through Yost Labs (https://yostlabs.com/3-space-application-programming-interface/). Python v2.7 was used to generate sounds, run the experiment and record data on a Dell Optiplex 725 intel Core 2 duo PC running Windows Vista. Participants responded via a keyboard by pressing the right or left arrow key and these responses were recorded using a custom-made python script.

### 2.3 Stimuli

Active head movement was self-generated by participants at the offset of a low pitch 200 Hz tone ‘go signal’ presented via headphones (Apple iPod earphones: MA662G/A). The duration of the go-signal ranged from 1-3s. The sound stimulus was a higher pitch 2000Hz tone presented for 50ms at a randomly generated time between 0 and 650ms after the offset of the go signal. At the beginning of the study, participants were seated in a chair, blindfolded with their eyes closed, and instructed to practice rotating their head to the right and then back before the actual trials commenced (c.f. [16]).

### 2.4 Procedure

Participants performed a temporal order judgement task in which they reported whether the onset of their head movement came first, or the onset of the high pitch sound stimulus came first. Each trial began with the onset of the low pitch go signal. The duration of the go signal was randomized to prevent participants from predicting the timing of the offset, and anticipating the start of the head movement ([c.f. [1]). At the offset of the go signal, participants initiated head movement, and due to the response time delay between the offset of the go signal and the onset of the head movement, the comparison sound stimulus could occur before or after the head movement. Participants responded by pressing the left or right key on the computer keyboard, where the left key indicated that the onset of head movement came first, and the right key indicated that the onset of the sound came first. Once the participant selected a response, the next trial would begin immediately after. A schematic of a typical trial is shown in Figure 1.

**Figure 1.**
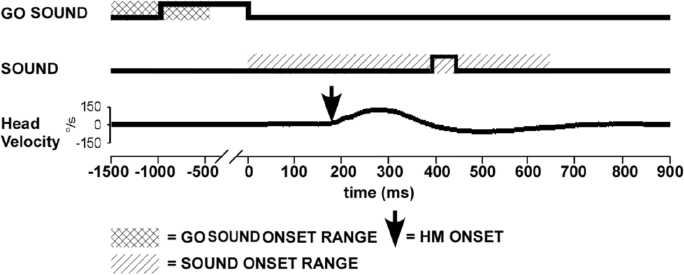
Example of a trial. Offset of the go sound is the signal to begin head movement (HM). The comparison sound is randomly generated between 0-650 ms after the go sound offset. Head movement can occur before or after the comparison sound stimulus due to the time it takes for the participant to perceive the go sound offset and initiate head movement.

Participants performed 10 practice trials prior to the experiment which then consisted of three conditions in a block design with 100 trials within each block. Each block took approximately 10 minutes to complete with a break of 5 minutes in between blocks. For the three conditions, participants were asked to move their head at what they subjectively considered to be a slow, normal, or fast head movement, the latter being as fast as they could move their head. The order of the conditions across participants was randomized.

### 2.5 Data Analysis

Raw recorded data was analyzed using Python 2.7. Angular velocity, which was originally recorded in raw form by the gyroscope in YEI 3-Space Data-logging sensor, was converted to degrees by accounting for the sensitivity level of 0.07º/sec/digit for ±500º/sec. Displacement was obtained using the formula:

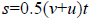

where *s* is the final displacement, *v* is the final velocity, *u* is the initial velocity, and *t* is the time at the final velocity. Angular acceleration was calculated by taking the change in velocity over the change in time between two adjacent data points. Onset of head movement was calculated to be 5ms before the velocity of the head was greater than three standard deviations from the average head velocity sampled 100ms before the trial onset (Figure 2). Each individual trial was further examined visually by plotting the angular velocity signal using the MatPlotLib library in Python 2.7. In trials where the onset of head movement was not accurately determined by the algorithm due to local minima or a noisy signal, the trial was discarded. Trials which had an excessively noisy signal or signals which had multiple peaks were removed from analysis. If greater than 20% of trials were removed for a condition for a participant, the participant was removed from analysis.

**Figure 2.**
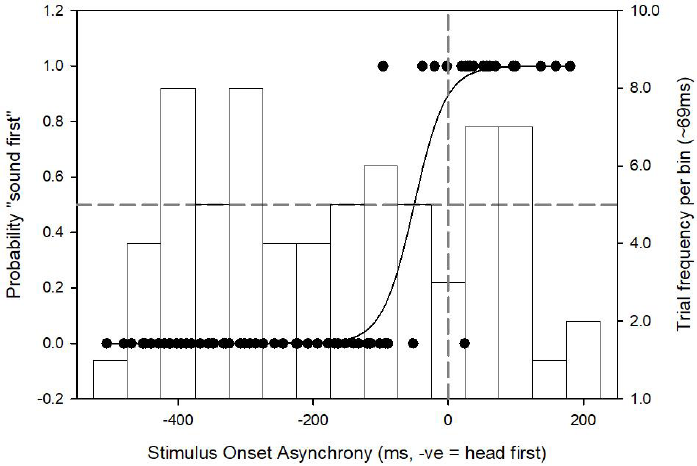
Sample TOJ data from an average participant from the fastest condition. Positive SOAs represent sound occurring first, whereas negative SOAs represent head movement occurring first. An SOA of 0ms represents true simultaneity and is represented by the dashed vertical line. The PSS occurs at a probability of 0.5 and is represented by a dashed horizontal line. A frequency distribution of SOAs binned for each 10% of trials is also shown on the right y-axis.

Due to the subjective nature of the participants deciding what constitutes a slow, medium and fast head movement and participants poorly replicating their head movement trajectory trial-to-trial [16], there was significant overlap in the peak velocities for the three conditions. To correct for this, the peak velocities of each participants were artificially stratified into four equally-sized conditions according to increasing peak velocity and renamed velocity 1, 2, 3 and 4. By stratifying into four, and not three conditions, more accurate linear regressions for within-subjects data could be obtained.

Stimulus onset asynchronies (SOAs) were determined by calculating the difference between head movement onset and sound onset, with a negative SOA indicating that the head moved prior to the sound. A logistic function (Eq. 2) was fitted to the participants responses for all four conditions as a function of SOA using SigmaPlot 12.5, with the inflection points of the logistic function (*x_0_*) taken as the point of subjective simultaneity (PSS; Figure 2) and the slope of the function (*b*) as the just noticeable difference (JND; [1]).

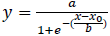

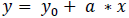

For the head movement dynamics, a one-way repeated-measures ANOVA was applied to peak velocity and time to peak velocity between the conditions to confirm that the different head movement categories were sufficiently significantly different from each other. One sample t-tests, or the Wilcoxon signed rank t-tests if data was not normally distributed as per the Shapiro-Wilk test, were conducted to compare the PSS of each condition to 0ms to confirm whether the head movement must precede sound stimulus to be perceived as simultaneous. To test whether there was a significant difference in PSS between conditions, a one-way repeated measures ANOVA (Holm-Sidak) was conducted between all four striated conditions. To assess the hypothesis that people who move their head faster require active head movement onset to occur earlier than a comparison sound stimulus (i.e., replicate [1]), we ran Pearson’s *r* correlations (Spearman’s *ρ* if not normally distributed) between peak head movement velocity and the PSS for each head movement condition, where a significant negative correlation for any head movement condition would replicate [1]. Lastly, to assess the hypothesis that the faster the head moves within-subjects requires active head movement onset to occur earlier than a comparison sound stimulus, a linear regression (Eq. 3) was fitted to each participant’s PSS values for each of the four velocity conditions, and an average linear regression line was obtained by taking the average of the slope (*a*) and y intercept parameters (*y*_0_) for the individual regressions. A one-sample t-test, or a Wilcoxon signed rank t-test if data was not normally distributed as per the Shapiro-Wilk test, of the average slope (*a*) relative to 0 (i.e., no change in the PSS relative to peak head movement velocity) would confirm this hypothesis if the average slope was negative, or the alternative hypothesis that an increase in stimulus intensity reduces the PSS if the average slope was positive. Tukey’s post hoc analysis was used to assess significant differences in all pairwise comparisons.

## 3. Results

Four artificial, equal-sized conditions were created by sorting the peak velocity of each participants from the lowest to highest velocity and then grouping the trials into four equally-sized conditions. These conditions are referred to as Velocity 1 (average: 76.46°/s, s.e.=6.42), Velocity 2 (average: 110.42°/s, s.e.=8.13), Velocity 3 (average: 167.47°/s, s.e.=12.30), and Velocity 4 (average: 256.78°/s, s.e.=19.75). In total, 6.47% of trials were removed due to anticipatory head movement, excessively noisy data, or two peaks being present in the velocity signal. Three participants were fully removed from the analysis since they did not contain one or more of the original three velocity conditions, due to excessively noisy trials, and could therefore not be striated into the four new conditions.

The mean and standard error of each head movement dynamics for every condition is shown in Figure 3. There were significant differences in peak velocity, time to peak velocity and time to peak displacement, confirming that participants indeed moved their head at different velocities. Post hoc analysis using Tukey’s procedure showed significant difference between the 1^st^ and 3^rd^, 1^st^ and 4^th^, and 2^nd^ and 4^th^ condition for both peak velocity and time to peak velocity. Given that varying the velocity of the head movement will affect the timing of the peak of the head movement dynamics, these results are expected.

**Figure 3.**
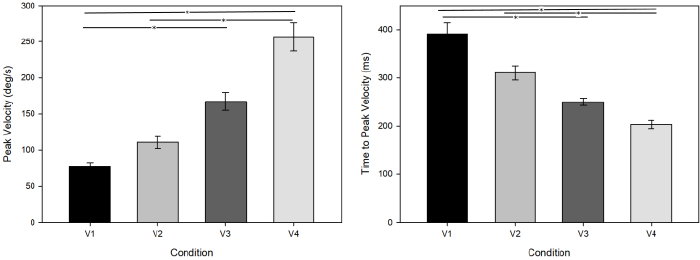
Average head movement dynamics for each stratified condition **a.** average peak velocity, **b.** average time to reach peak velocity. Error bars are ±1 SEM. *: p < 0.05

Figure 4a-d shows the results of fitting the logistic curve function to each individual participant’s data (grey lines and dots) as well as a representation of the average logistic curve constructed from the average slope and PSS value for each condition (black lines and dots). Figure 4e shows the individual (grey dots) and average (black dot with standard error bars) PSS values for each condition. In the Velocity 1 condition, the average PSS was -122.51 ms (s.e.=18.32) and significantly before 0ms (t(16)=-6.688, p<0.001). In the Velocity 2 condition, the average PSS was -110.94 ms (s.e.=20.60) and significantly before 0ms (Median=-103.5243, Wilcoxon Z=-3.621, p<0.001). In the Velocity 3 condition, the average PSS was -66.57 ms (s.e.=22.19) and significantly before 0ms (t(16)=-3.000, p=0.00848). In the Velocity 4 condition, the average PSS was -52.13 ms (s.e.=22.76) and significantly before 0ms (t(16)=-2.290, p=0.0359). The global average PSS value for all four conditions was -88.03 ms (s.e.=10.89). There was no significant difference for the JND values between the four conditions (Figure 4f), meaning that the participants' precision did not differ as the velocity of head movement changed. Together these results support our first hypothesis, and replicate previous work showing that the perceived timing of an active head movement is slow compared to a comparison sound stimulus [1,16,17].

**Figure 4.**
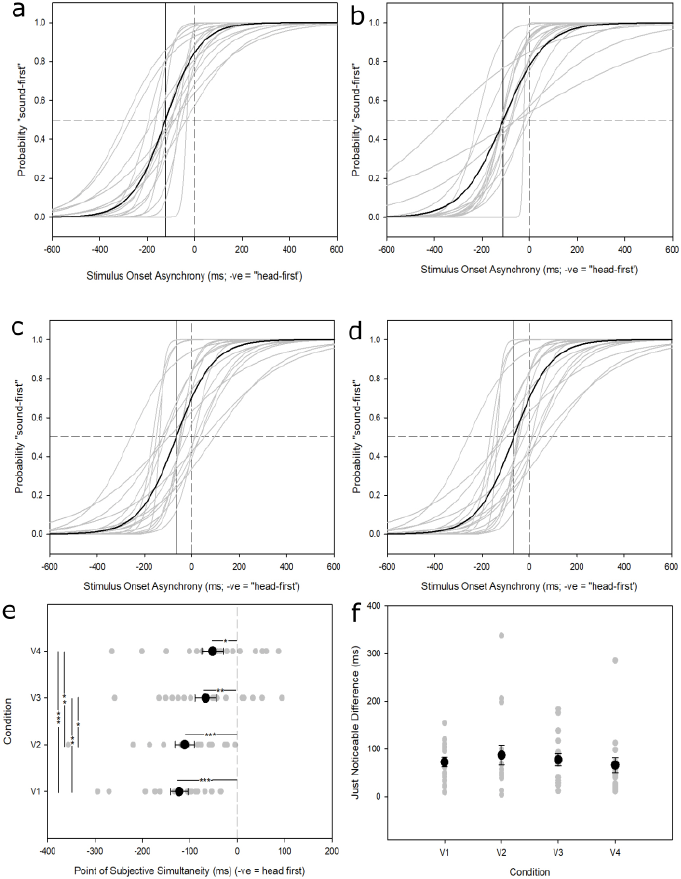
Average TOJ, PSS and JND data for all four stratified velocities. **a**. Slowest velocity (V1) TOJ data, with gray curves representing individual participants, and the black curve representing the average logistic function. **b**. Second-slowest (V2) TOJ data. **c.**, Second-fastest (V3) TOJ data. **d**, Fastest (V4) TOJ data. **e**. Average PSS data for all four stratified conditions. Grey dots represent individual participants and black dots represents the average PSS for each condition, with standard error bars. **f**. Average JND data for all four stratified conditions. Grey dots represent individual participants and black dots represent average JND value for each condition. Error bars are ±1 SEM. *: p<0.05, **: p<0.01; ***: p<0.001.

A one-way repeated measures ANOVA (Holm-Sidak) indicated a significant difference in PSS values between subjects (F(3,67)=9.39, p<0.001) and Tukey’s post hoc analysis showed significant differences for all pairwise comparisons (p<0.05; Figure 4e) except between Velocity 1 and Velocity 2 (p=0.464) and between Velocity 3 and Velocity 4 (p=0.593). Correlations between peak velocity and time to peak velocity versus PSS were run separately for each velocity condition. Peak velocity had no significant relationship to the PSS for Velocity 1 (Pearson’s r=0.157, p=0.548), Velocity 2 (Spearman’s *ρ*=0.061, p=0.817), Velocity 3 (Pearson’s r=0.086, p=0.741), or Velocity 4 (Pearson’s r=0.068, p=0.794). Neither did the time to peak velocity versus have any significant relationship to the PSS for Velocity 1 (Spearman’s *ρ*=0.191, p=0.461), Velocity 2 (Spearman’s *ρ*=0.123, p=0.639), Velocity 3 (Pearson’s r=0.256, p=0.321), or Velocity 4 (Pearson’s r=0.325, p=0.203) suggesting that the speed of the active head movement does not have an influence on the PSS, which causes us to reject our second hypothesis.

To test our third hypothesis, linear regressions of peak velocity versus PSS were applied individually for each participant, and are shown in Figure 5a. From the slopes and intercepts of these linear regressions, an average regression line was obtained, to describe the overall trend within-subjects (Figure 5b and 5c). The average regression line has a slope of 0.682 (s.e.=0.3211; median=0.892. A one-sample signed-rank test confirmed that the regression slopes are significantly different from zero (Median=0.892, Wilcoxon Z=2.49, p=0.011). These results suggest that within-subjects, an increase in active head movement velocity leads to a smaller PSS and we reject our third hypothesis.

**Figure 5.**
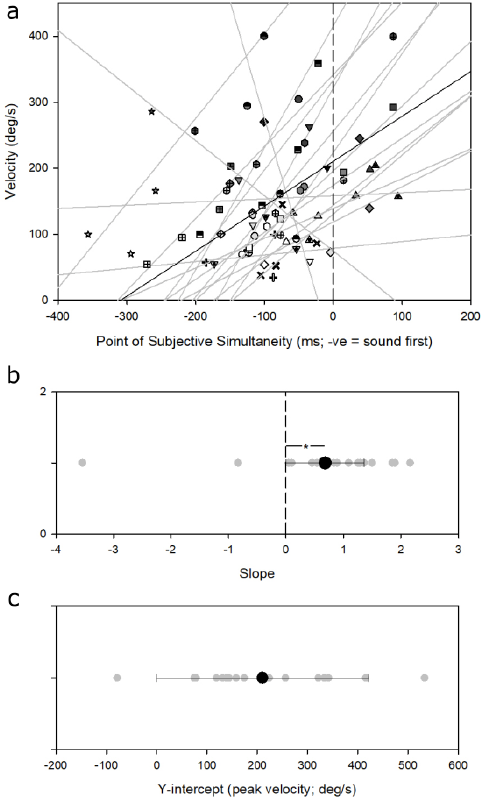
**a.** Individual linear regressions for each participant for all four velocity conditions. Different symbols represent different participants. Thicker black line represents the linear regression for the average participant. Dashed line shows the point of true simultaneity. **b.** Average Slope and **c.** Y-intercept for within-groups linear regressions. Each gray dot represents one participant, the black dot represents the average. Error bars are ±1 SEM. *: p<0.05.

## 4. Discussion

In the present study, we investigated whether the velocity of active head movement will influence the perceived timing of the head movement using a within-subjects design. We provide further evidence that the perceived timing of active head movements is slow when paired with a sound stimulus. This delay, which had a global average of -88 ms for all conditions is similar to the 80ms delay previously reported [1,16,17], although it is important to note that previous studies only looked at one active head movement velocity, whereas our study looked at a range of active head movement velocities. Contrary to the predictions of our second hypothesis, the results showed that a greater peak velocity did not lead to a significantly greater delay in the PSS when comparing among each individual group between-subjects. Most importantly, the individual regressions of the within-subjects data revealed that an increase in peak head movement velocity is significantly correlated with a reduction in the delay of the PSS.

Barnett-Cowan and Harris[1] reported an increased lag in the perceived timing of active head movements as the velocity of head movement increased, in a between-subjects design. We find no further evidence of this in our study and quite convincingly show that higher velocities cause a decrease in the lag of the perceived timing of an active head movement, and not an increase. It should also be noted that two other studies since [1] have also found no effect of head movement velocity on the PSS between-groups [16,17].

Our within-subjects result that the perceived timing of active head movements becomes less delayed at increasing head velocities are in agreement with other literature on stimulus intensity [9-15]. Here, the timing between stimuli to be perceived as simultaneous is shorter when the intensity is increased. A greater head movement velocity may be considered a more intense stimulus, as it requires the neck muscles to generate a larger force and evoking stronger sensory signals from the vestibular and neck proprioception neurons. This could also explain why we only observe a significant difference in the within-subjects data because we can only compare the varying intensity within individuals, due to the subjective nature of our stimuli. This further strengthens the hypothesis that the perceived timing of an active head movement can be modulated by the intensity of the stimuli, represented by the velocity of the head movement. One possible reason that the perceived timing of active head movements is more delayed at lower velocities is that the brain increases its tolerance for the asynchrony of a weaker stimulus[14].

It is known that when an efferent signal is sent from the motor cortex to the muscles, an efference copy, also known as corollary discharge, is created which gets routed to other parts of the sensory cortex [8]. This efference copy can then modify the excitability of other sensory areas. There are two competing hypotheses concerning the perceived timing of active head movements. The first, known as the anticipation hypothesis, postulates that the efference copy of the active head movement will allow head movement to be perceived quicker than a passive head movement. On the other hand, this efference copy could also be used to suppress the vestibular nucleus, which would delay the perceived onset time.

In the previous study by Barnett-Cowan and Harris [1], passive head movements were perceived earlier than active head movements. These results seemed to point towards the suppression hypothesis, where an efference copy of the active movement suppresses the vestibular nucleus and delays the perceived timing of the head movement. Additionally, it appeared as though the mechanism for the suppression could be velocity-based. A greater head movement velocity was correlated with a larger delay in PSS, similar to the findings of vestibular suppression of active head movements in monkeys [7]. If increasing speeds of active head movements increase the delay in perception, it would provide further evidence for velocity-based suppression. However, since we found that increasing the speed of active head movement decreases the delay in perception within-subjects, our results do not support a velocity-based suppression. Further studies should look at the effect of velocity with both active and passive head movement to determine whether the findings of Barnett-Cowan and Harris [1] can be replicated when explicitly controlling for the velocity of the head movement.

From Figure5a, we can see that the slopes cluster around 0-2, except for two participants, which have negative slopes. Given that most participants had slopes that were relatively close to one another, we suspect that the results from these two participants are not indicative of the typical participant. These negative slopes may be a result of the small number of data points that were used to make the linear regressions, a result of the constraints in the number of trials that could be conducted for each participant, and the minimum number of trials that were necessary to create the corresponding psychometric functions.

From the within-groups analysis, it is suggested that true simultaneity of audio-vestibular stimuli would be reached at around 200 °/s. However, it is important to note that the within-groups comparison only contained four data points per participant for each linear regression, for each of the four conditions. This limits any analysis on the dynamics that head movement velocity has on PSS. We cannot conclude whether the behavior is linear, or higher order, and importantly how these change across individuals. A visual analysis of the within-group regression seems to indicate a more exponential relationship. Future studies could include more trials per participant so that the velocities can be stratified into more than four conditions, to allow for a closer analysis of the dynamics of this effect and to tease apart whether this relationship is linear or higher order.

## 5. Conclusion

From the results of this experiment, we conclude that the perceived timing of active head movement is slow in comparison to an auditory stimulus, replicating previous research on the perceived timing of active head movements. Furthermore, we conclude that an increased active head movement velocity shortens this perceptual delay in a within-subjects design. This is in line with literature where more intense auditory, visual and vestibular stimuli are perceived earlier in time. We failed to replicate the between subjects results from Barnett-Cowan and Harris [1] where an increase in the velocity of active head movements led to an increase in the perceptual delay when paired with a comparison auditory stimulus. Although our results do not refute the suppression hypothesis that was previously reported, where an efference copy of the active head movement delays the perceived timing of the head movement via suppression of the vestibular afferent signals [1], we do provide evidence against a velocity-based suppression mechanism.

## Acknowledgments

Supported by an Natural Sciences and Engineering Research Council of Canada (NSERC) Discovery Grant (#RGPIN-05435-2014) to MB-C. We thank Nazanin Mohammadi for computer programming and helping to test participants.

## Conflict of interest

None.

